# Scaffolding protein IQ motif containing GTPase activating protein 2 (IQGAP2) regulates glycogen metabolism

**DOI:** 10.1101/2020.05.28.121632

**Authors:** Anushna Sen, Sara Youssef, Karen Wendt, Sayeepriyadarshini Anakk

## Abstract

The liver is critical in maintaining metabolic homeostasis, regulating both anabolic and catabolic processes of fats, proteins, and carbohydrates. The IQ motif containing GTPase activating protein 2 (IQGAP2) is a member of the IQGAP family. Of the three homologous isoforms, the IQGAP2 scaffolding protein is predominantly found in the liver. To characterize its role in regulating metabolism, *Iqgap2*^*−/−*^ female and male mice, and their WT controls, were fed ad libitum or fasted for 24 hours. Hepatic gene expression, protein levels, and the metabolic response were compared between WT and *Iqgap2*^*−/−*^ mice, using RT-qPCR, western blot analysis, and histological stains. We found that loss of IQGAP2 alters the phosphorylation of active glycogen synthase kinase 3 (GSK3) expression, a known regulator of glycogen synthesis and lipogenesis. Consistent with this result, *Iqgap2*^*−/−*^ female mice displayed depletion of periportal glycogen even in the fed state. We also observed the blunted expression of genes involved in glycogenesis and lipogenesis when IQGAP2 was deleted. Since GSK3 is known to regulate the activity of β-catenin, we examined and found it to be reduced in *Iqgap2*^*−/−*^ mice. Our findings demonstrate that IQGAP2 plays an important role in regulating glycogen synthesis.

## Introduction

The liver tightly regulates lipid and glucose metabolism to maintain energy homeostasis (1, 2). Hepatocytes are thus programmed to modulate metabolism and adapt to nutrient fluctuations in fed and fasted conditions. During the fed state, glycolysis, glycogenesis, and *de novo* lipogenesis are all induced in the liver. GSK3, a crucial regulator of glycogen synthesis inactivates glycogen synthase (GYS2) and inhibits glycogenesis (1, 3). Phosphorylation of GSK3 inactivates the kinase and thus leads to GYS2 activation in the fed state. Conversely, dephosphorylation of GSK3 causes GYS2 phosphorylation thus inactivating GYS2. In this way, GSK3 controls glycogen synthesis (1, 3). In addition, in nutrient excess, *Srebp1c* regulates fatty acid synthase (*Fasn*) to promote *de novo* lipogenesis (3, 4). Under normal conditions, only a small quantity of fatty acids is stored as triglycerides, with diacylglycerol acyltransferases (*Dgat2*) catalyzing the final step of triglyceride synthesis (5). Under fasting conditions breakdown of glycogen, gluconeogenesis, and fatty acid oxidation are upregulated (2, 3). As glycogen stores are depleted, the primary fuel source shifts from carbohydrate-to fat-based (6, 7). Fat is broken down into fatty acids, through fatty acid oxidation, with carnitine palmitoyltransferase-1a (*Cpt1a*) serving as a rate-limiting step (7–9). This process helps to generate ATP, providing fuel for the cells during prolonged fasted state (3).

IQGAP2 is a Ras GTPase-activating-like protein (10), which binds to activated CDC42 and RAC1, but it does not stimulate their GTPase activity (10–13). The IQGAP family consists of three highly homologous members (IQGAP1, IQGAP2, and IQGAP3), with IQGAP1 being the most studied isoform (14–21). While IQGAP1 is ubiquitously expressed, IQGAP2 is mainly expressed in the liver and platelets (22). Previous data revealed a potential role for IQGAP2 in regulating hepatic fatty acid uptake and lipid processing (23, 24). We recently showed that IQGAP1 was crucial for regulating ketogenesis under fasting conditions (20), therefore, we investigated the role for IQGAP2 in maintaining energy metabolism. First, we characterized the *Iqgap2*^*−/−*^ female and male mice in fed and fasted conditions. We examined lipid and carbohydrate metabolism and identified a role for IQGAP2 in regulating glycogen levels.

## Results

### IQGAP2 females are resistant to fasting-induced weight loss

Both sexes of WT mice lost weight upon fasting (Table 1), *Iqgap2*^*−/−*^ males mimicked the same trend whereas *Iqgap2*^*−/−*^ females did not lose weight. Reduction in gonadal weight was noted only in the WT female mice. However, fasting-induced decrease in liver weight was observed in all the groups of mice except *Iqgap2*^*−/−*^ females. (Table 1). Next, we examined if the loss of *Iqgap2* was compensated by overexpression of *Iqgap1* and/or *Iqgap3.* We examined their transcript levels and found no significant differences between WT and *Iqgap2*^*−/−*^ mice (Fig. S1). We also validated this result and found no changes in IQGAP1 protein levels when IQGAP2 was deleted (Fig. S1). Since we saw changes in liver weight with fasting, we characterized and compared liver histology of WT and *Iqgap2*^*−/−*^ mice using hematoxylin and eosin staining. No difference in liver histology was observed either in IQGAP2 deficiency (Fig. S2) or between the fed and fasted states. These results revealed loss of IQGAP2 does affect the liver architecture but alters fasting-induced weight loss in females.

**Table 1.** Gross organ and body weights of fed and 24hour fasted wild-type and *Iqgap2*^*−/−*^ mice (male and female, 14-16wks old, n=5-15).

| **Gross Morphology** | **Wild-type Fed** | **Wild-type Fasted** | ***Iqgap2*<sup>-/-</sup> Fed** | ***Iqgap2*<sup>-/-</sup> fasted** |
| --- | --- | --- | --- | --- |
| *Males* |  |  |  |  |
| Body weight (g) | 27.10 ± 2.76 | 23.72 ± 4.5* | 29.48 ± 3.47 | 25.51 ± 2.84* |
| Liver weight (g) | 1.01 ± 0.25 | 0.75 ± 0.08* | 1.02 ± 0.14 | 0.72 ± 0.13*** |
| WAT weight (g) | 0.76 ± 0.28 | 0.61 ± 0.21 | 0.80 ± 0.26 | 0.71 ± 0.38 |
| *Females* |  |  |  |  |
| Body weight (g) | 21.51 ± 2.91 | 18.42 ± 2.88* | 24.76 ± 3.85 | 24.44 ± 4.15 |
| Liver weight (g) | 0.85 ± 0.16 | 0.68 ± 0.12* | 0.83 ± 0.22 | 0.71 ± 0.15 |
| WAT weight (g) | 0.69 ± 0.26 | 0.39 ± 0.21* | 0.49 ± 0.23 | 0.62 ± 0.34 |

Statistics were calculated using unpaired t-test. \*p<0.05 and \*\*\*P<0.001 vs fed control.

### IQGAP2 knockout livers revealed a blunted lipid metabolic response

Since lipid levels are altered between the fed and fasted states, we explored if IQGAP2 functioned to regulate this process. In the fed state, the liver upregulates lipogenic processes. Both *Iqgap2*^*−/−*^ females and males displayed reduced expression of *Srebp1c* (Fig. 1A, E), a key transcription regulator of *de novo* lipogenesis. Concomitantly, *Iqgap2*^*−/−*^ females displayed significant decreases in the expression of lipid synthesis genes *Fasn* and *Dgat2* compared to WT livers (Fig. 1B, C). This genotypic reduction in gene expression was unique to females, as *Iqgap2*^*−/−*^ male mice showed comparable expression profile to that of male WT controls (Fig. 1F, G). Additionally, this blunted lipid metabolic response was also evident in the fasted condition in *Iqgap2*^*−/−*^ female mice. Typically, liver upregulates fatty acid oxidation by increased gene expression of the rate-limiting enzyme CPT1a (7–9), and this effect was impaired in *Iqgap2*^*−/−*^ fasted female but not in male mice compared to their respective WT control (Fig. 1D, 1H). This data indicates that IQGAP2 deficiency alters hepatic lipid metabolism in a sex-specific manner. To see if this effect was due to sexually dimorphic expression of IQGAP2, mRNA and protein levels were measured in WT fed and fasted control mice in both sexes (Fig. S3). Despite significantly lower *Iqgap2* transcript expression, protein levels were comparable in both sexes. Overall, our data suggest that female *Iqgap2*^*−/−*^ livers have impaired lipid metabolism.

**Fig. 1.**
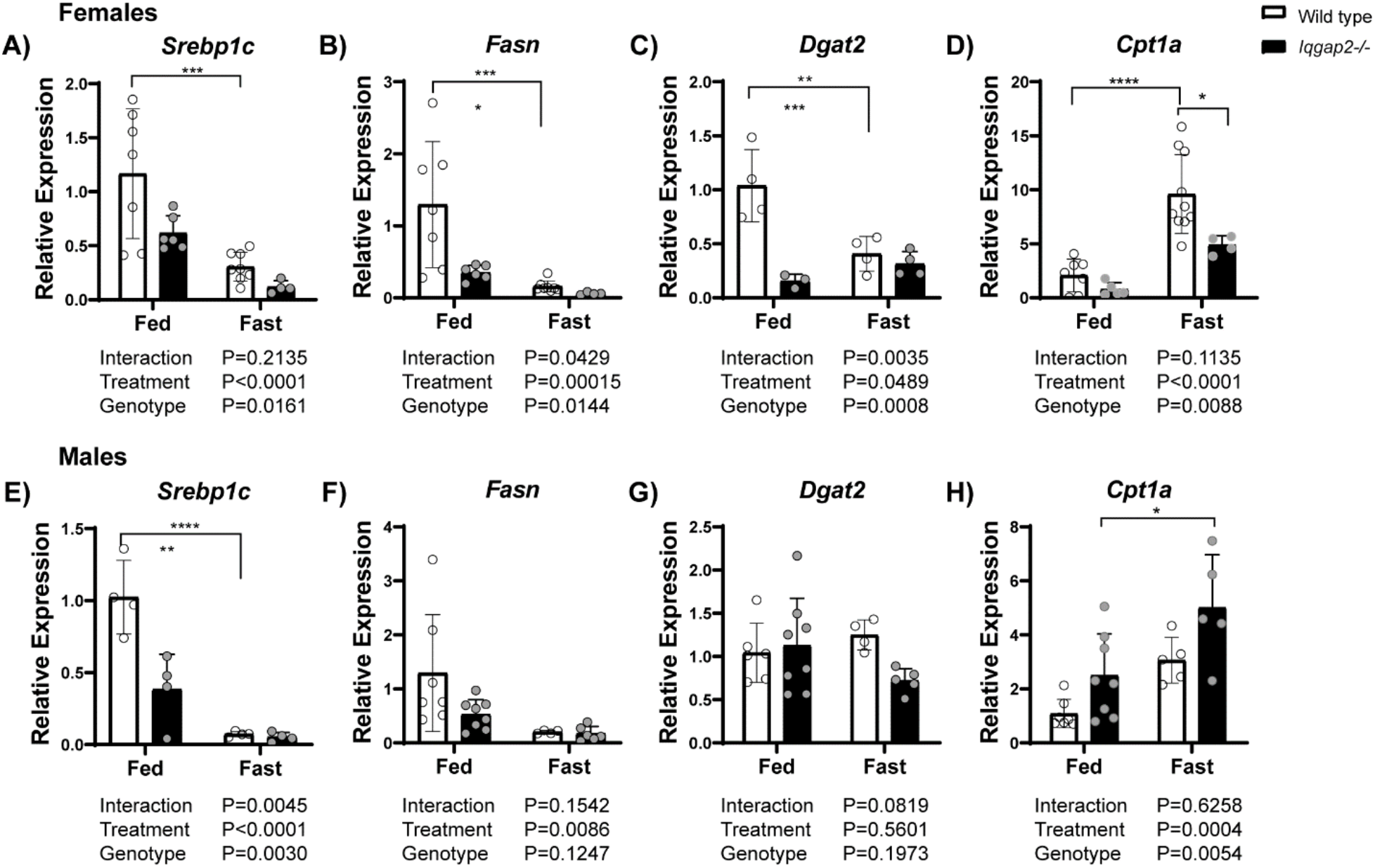
Depletion of Iqgap2 blunts hepatic lipid metabolism in female mice. *Iqgap2*^*−/−*^ and wild-type mice (male and female, 14-16 weeks old, n=5-15) were fed regular chow ad libitum or fasted for 24 hours. Liver tissues were analyzed with qRT-PCR to quantify mRNA of key genes involved in lipid metabolism. Lipogenic transcription factor *Srebp1c* was downregulated in both female and male *Iqgap2*^*−/−*^ mice (A,E). However, (B) *Fasn* and (C) *Dgat2* were reduced in female but not in *Iqgap2*^*−/−*^ male mice (F, G). Fasting mediated increases in *expression of Cpt1a gene, crucial in* fatty acid oxidation *was diminished in f*emale *Iqgap2*^*−/−*^ mice (D). In contrast, male mice showed significant upregulation *of Cpt1a* upon fasting (H). Statistics were calculated using two-way ANOVA with Bonferroni post hoc analysis. *P<0.05, **P<0.01, ****P<0.0001.

### Female *Iqgap2*^*−/−*^ mice displayed an altered carbohydrate metabolism in fed state

We then evaluated whether loss of IQGAP2 affects other metabolic processes such as carbohydrate metabolism. We analyzed transcript levels of genes regulating glycogenesis (*Gys2*), gluconeogenesis (glucose 6-phosphatase, *G6pase*, and fructose 1,6-bisphosphatase, *Fbpase*), and glycolysis (phosphofructokinase, *Pfkl*, and glucose transporters 1 and 2, *Glut1* and *Glut2*, respectively) (25–28). *Iqgap2*^*−/−*^ female mice displayed compromised carbohydrate metabolism in the fed state. *Iqgap2*^*−/−*^ female livers displayed a five-fold decrease in *Gys2* mRNA levels (Fig. 2A). In contrast, *Iqgap2*^*−/−*^ male mice displayed a two-fold increase in *Gys2* compared to their WT controls (Fig. 2D). Further, we observed a decreasing expression pattern of gluconeogenic genes, with two-fold lower *Fbpase* and *G6pase* in *Iqgap2*^*−/−*^ females (Fig. 2B, C) but not in the male mice (Fig. 2E, F). Additionally, genes involved in glycolysis and glucose transport remained unaltered between *Iqgap2*^*−/−*^ and WT mice (Fig. S3). Since reduced expression of both lipogenic and carbohydrate metabolic genes was distinctly observed in female *Iqgap2*^*−/−*^ mice, not males, we focused on the female mice for further analysis.

**Fig. 2.**
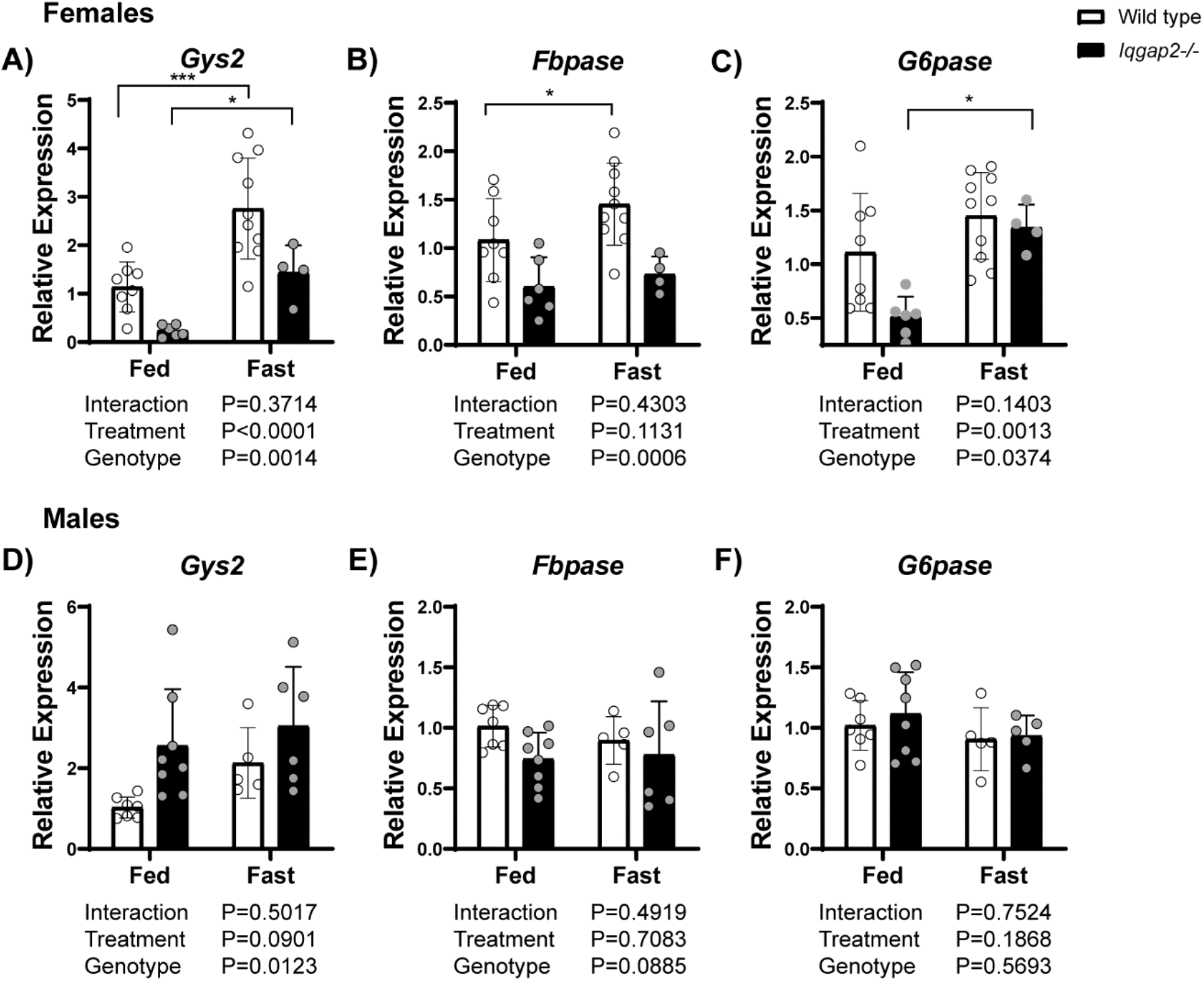
Carbohydrate metabolism is altered in female Iqgap2^−/−^ mice. Liver tissues from *Iqgap2*^*−/−*^ and wild-type mice were examined to quantify expression of genes involved in glycogen synthesis and gluconeogenesis. Glycogen synthetase gene expression was differential between the two sexes in *Iqgap2*^*−/−*^ mice (A, D). Female *Iqgap2*^*−/−*^ mice also showed blunted gluconeogenic response upon fasting with decreased expression of (B) *Fbpase.* (C) *G6pase* expression was lower in female mice in fed state but fasted response was comparable wild-type controls. Male *Iqgap2*^*−/−*^ mice do not show any deficit and showed expression pattern comparable to their wild-type controls (E, F). Statistics were calculated using two-way ANOVA with Bonferroni post hoc analysis. *P<0.05, ***P<0.001.

### Genes regulating nutrient signal and mitochondrial complex are intact in *Iqgap2*^*−/−*^ female livers

Since fibroblast growth factor receptor 1 (*Fgfr1*), fibroblast growth factor co-receptor *β-Klotho*, and fibroblast growth factor 21 (*Fgf21*) coordinate glucose and fat homeostasis (29–32), we examined their expression. Transcriptional levels of *Fgfr1, β-Klotho*, and *Fgf21* remained comparable in *Iqgap2*^*−/−*^ female mice to their WT control (Fig. 3A-C). Further, the fasting-induced *Fgf21* response (Fig. 3C) was maintained in *Iqgap2*^*−/−*^ mice. Since mitochondrial function is central to maintaining metabolic processes, we next analyzed the effects of IQGAP2 deficiency on the mitochondria. Transcript levels of peroxisome proliferator-activated receptor-γ coactivator (*Pgc1-a*), which regulates mitochondrial biogenesis (33, 34), and the mitochondrial cytochrome c oxidase subunits II and III (*CoxII* and *CoxIII*) were examined (Fig. 3D-F). Expression of these transcripts was intact in *Iqgap2*^*−/−*^ and similar to that of WT mice. This data suggests that the defective fed state response in *Iqgap2*^*−/−*^ female mice is not secondary to poor mitochondrial function.

**Fig. 3.**
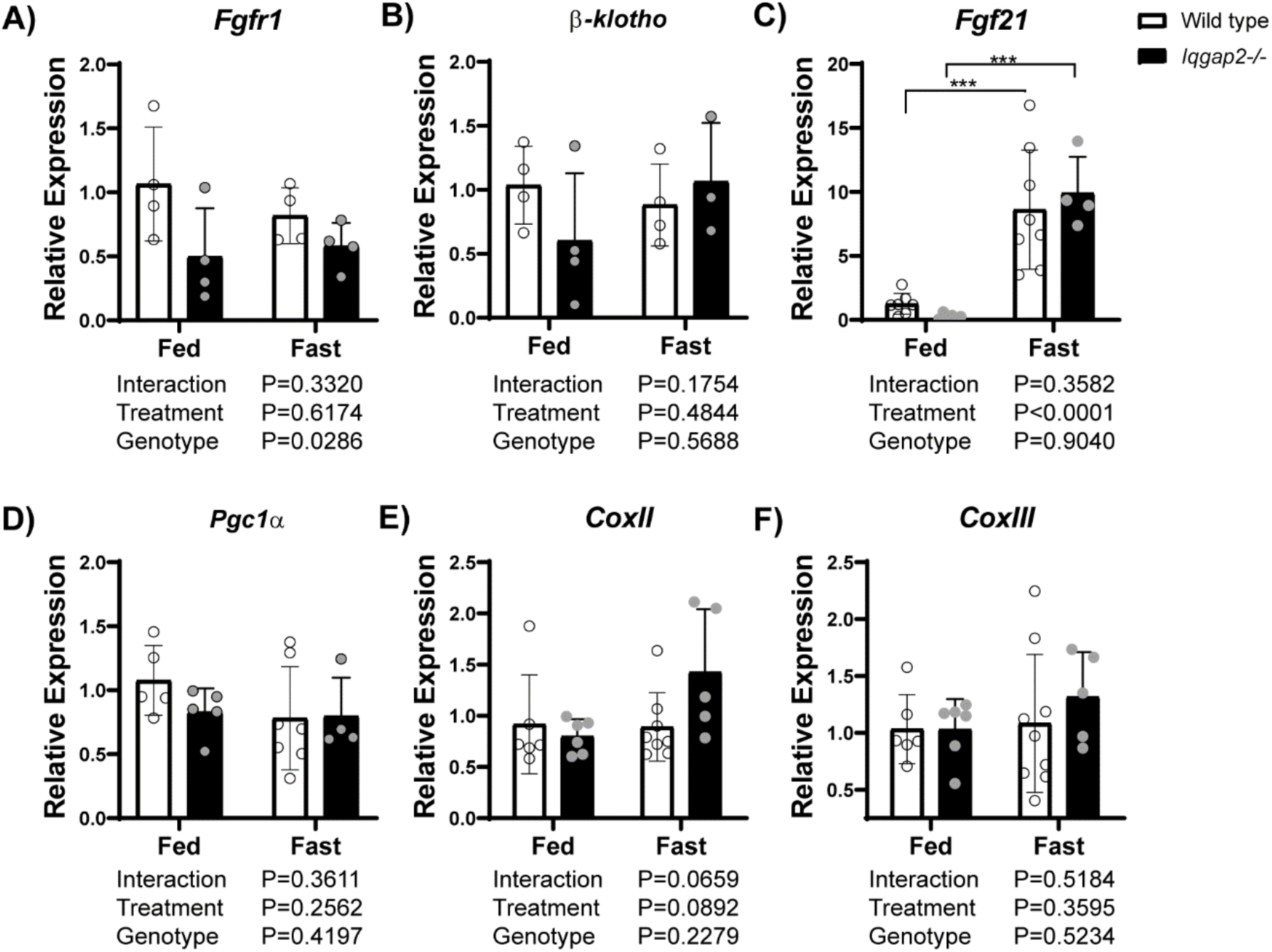
Iqgap2^−/−^ livers do not exhibit defects in nutrient signals and mitochondrial function. Enterokine receptor (A) *Fgfr1*, and coreceptor (B) *β-klotho* as well as fasting response (C) *Fgf21* gene expression was quantified using qRT-PCR. Fed state levels were comparable and the fasting-mediated increase in Fgf21 was unaltered in *Iqgap2*^*−/−*^ mice. Transcript levels of the master regulator of mitochondrial biogenesis (D) *Pgc1-α* and mitochondrial complexes II (E) and III (F) were similar between the groups. Statistics were calculated using two-way ANOVA with Bonferroni post hoc analysis. N=4-15 mice per group. ***P<0.001.

### Depletion of hepatic glycogen is observed in *Iqgap2*^*−/−*^ female livers

Due to the defects seen in hepatic lipid and carbohydrate metabolism in fed states, we explored if glycogen synthesis is defective in IQGAP2 deficient mice. Since *Gys2* gene levels (Fig. 2A) were lower in *Iqgap2*^*−/−*^ fed mice, we examined the functional significance of it. We stained WT and *Iqgap2*^*−/−*^ liver samples with PAS to analyze glycogen levels (Fig. 4). Hepatic glycogen was uniformly distributed in WT fed mice; however, glycogen staining was significantly depleted around the periportal veins in *Iqgap2*^*−/−*^ mice in the fed state. Since PAS stains both glycogen and glycoproteins, we investigated the periportal loss of PAS staining in the presence and absence of diastase (Fig.4), which digests glycogen. In the presence of diastase, we did not see any difference between WT and *Iqgap2*^*−/−*^ livers indicating that the loss of glycogen was responsible for reduced staining in them. Next, we examined and found fasting-induced marked loss of glycogen was similar between WT and *Iqgap2*^*−/−*^ mice, suggesting that mobilization of glycogen remained intact in the absence of IQGAP2 (Fig. S4). To see if a similar glycogen defect was displayed in *Iqgap2*^*−/−*^ fed male mice, WT and *Iqgap2*^*−/−*^ male liver samples were also analyzed using PAS (Fig. S5). We did observe a subtle decrease of glycogen levels around the periportal veins in *Iqgap2*^*−/−*^ fed male mice even though the gene expression of *Gys2* was not different. Similar to females, fasting hepatic glycogen levels were unaltered between *Iqgap2*^*−/−*^ and WT control liver. Our data displays a periportal glycogen depletion in *Iqgap2*^*−/−*^ fed mice that is exaggerated in female livers. This demonstrates that IQGAP2 plays a role in glycogen synthesis in the fed state.

**Fig. 4.**
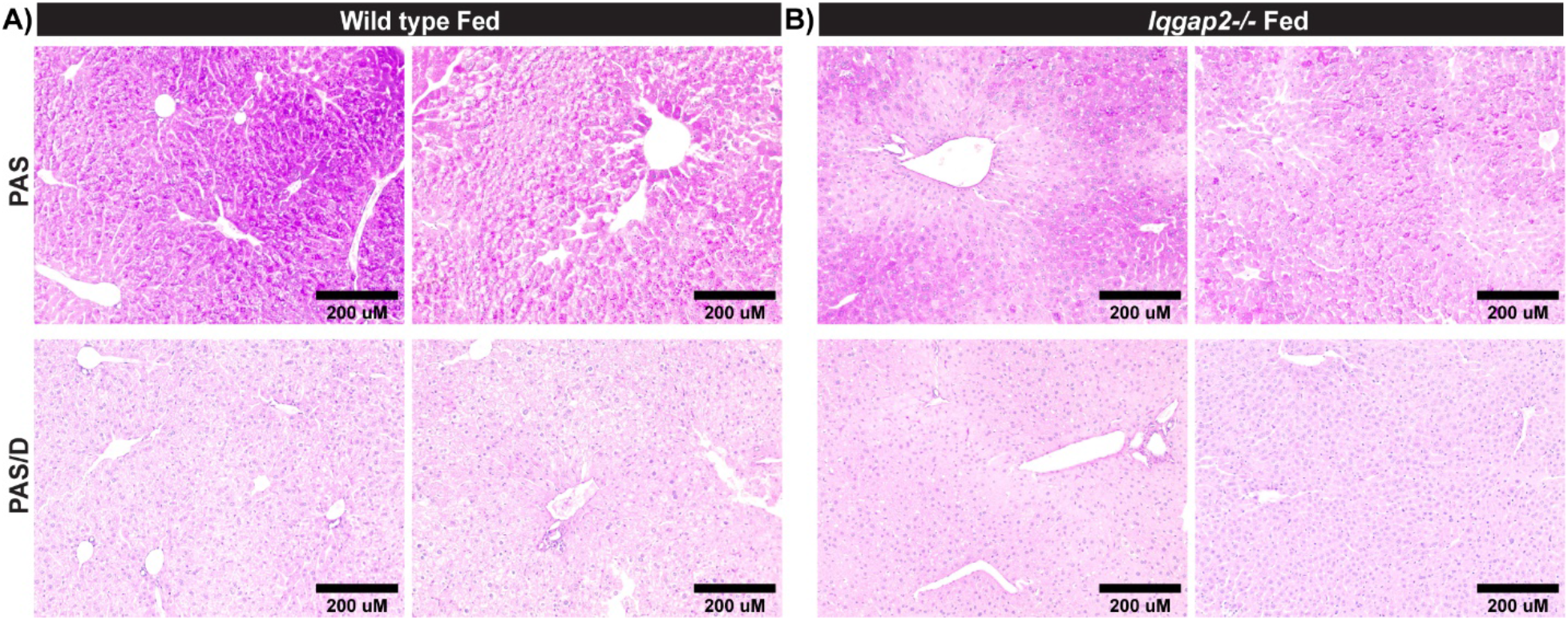
Significant reduction in glycogen storage occurs in Iqgap2^−/−^ female livers. *Iqgap2*^*−/−*^ and WT mice were fed regular chow ad libitum. Liver tissue sections were analyzed for glycogen levels by Periodic Acid-Schiff stain (PAS). Histological examination demonstrates uniform distribution of hepatic glycogen in the fed state of (A) wild-type mice whereas periportal depletion of glycogen is apparent in (B) *Iqgap2*^*−/−*^ mice. Each panel represents an individual mouse. Diastase treatment digests glycogen, so PAS/D+PAS is used to examine glycogen levels. N=3-4 mice per group. Bar, 200μM.

### IQGAP2 modulates GSK-3 activity

Because IQGAP2 deletion reduced glycogen levels, we analyzed GSK3 activity. Glycogen synthase is active when GSK3 is inactivated by phosphorylation (3). Conversely, unphosphorylated GSK3 can phosphorylate glycogen synthase to inactivate it. Therefore, we performed immunoblots and analyzed total and p-GSK3 levels (Fig. 5A). We found that the ratio of p-GSK3 to GSK3 was robustly reduced in *Iqgap2*^*−/−*^ fed mice compared to WT fed mice (Fig. 5B). This reduction suggests that inhibition of hepatic glycogen synthase occurs in the absence of IQGAP2.

**Fig. 5.**
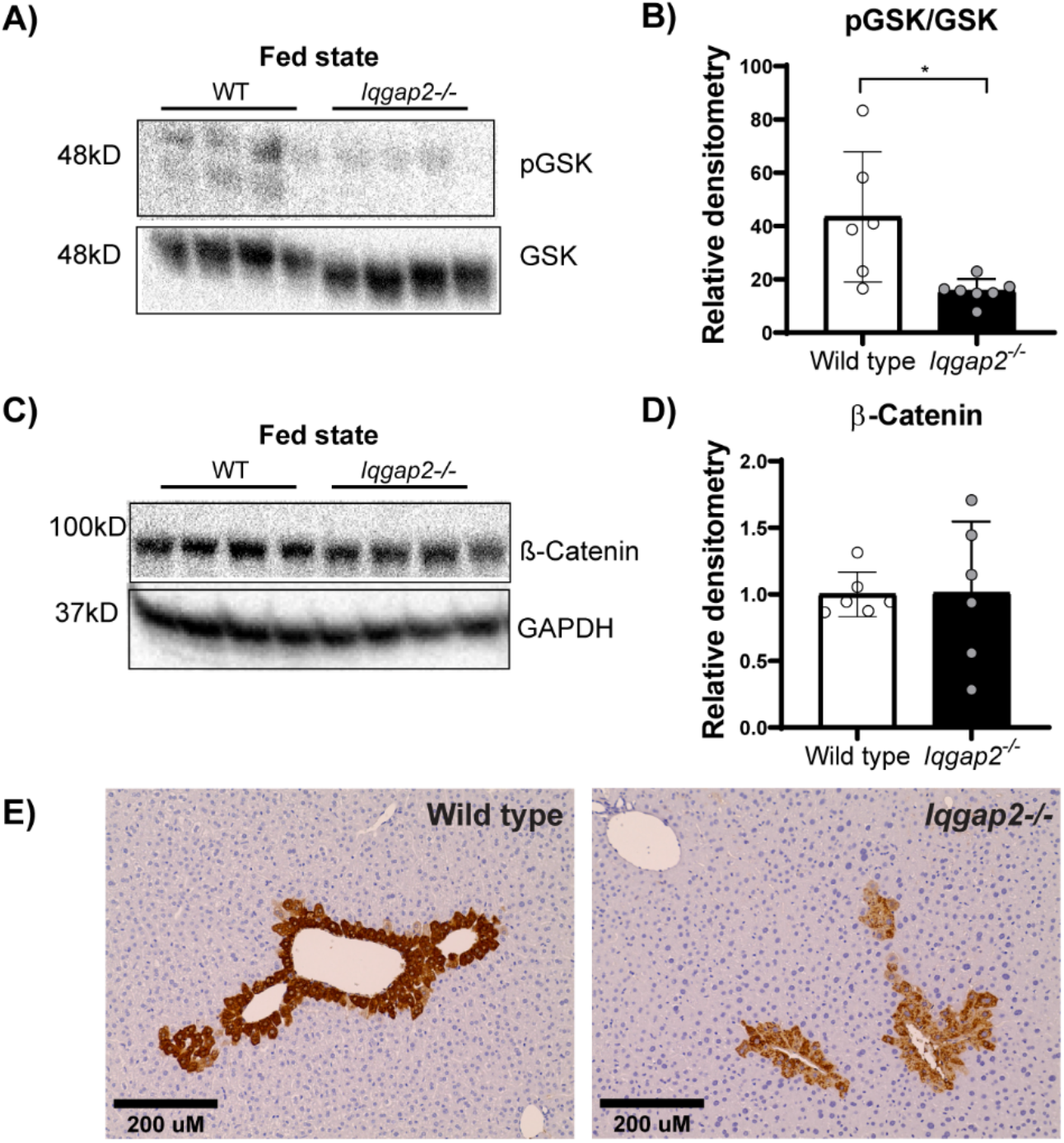
Iqgap2 depletion results in reduced inactivation of GSK3 signaling. Liver tissues from *Iqgap2*^*−/−*^ and wild-type mice were examined by western blot and immunohistochemical analysis. Hepatic (A) total GSK3β and phosphorylated GSK3β (ser9) protein levels were measured by western blots. Densitometric analysis of (B) inactive GSK3β (ratio of pGSK to GSK) indicated increased GSK3β activity in *Iqgap2*^*−/−*^ mice. To validate this increased GSK3β activity, we investigated its downstream target β-catenin in *Iqgap2*^*−/−*^. (C, D) β-Catenin protein levels were unaltered between *Iqgap2*^*−/−*^ and wild type mice. However, when we examined β-Catenin activity by measuring its key target glutamine synthetase, we found it decreased in *Iqgap2*^*−/−*^ mice (E). Glyceraldehyde-3-phosphate dehydrogenase (GAPDH) served as loading control for immunoblot analysis. Statistics were calculated using two-way ANOVA with Bonferroni post hoc analysis. *P<0.05. N=3-4 mice per group. Bar, 200μM.

**Fig. 6.**
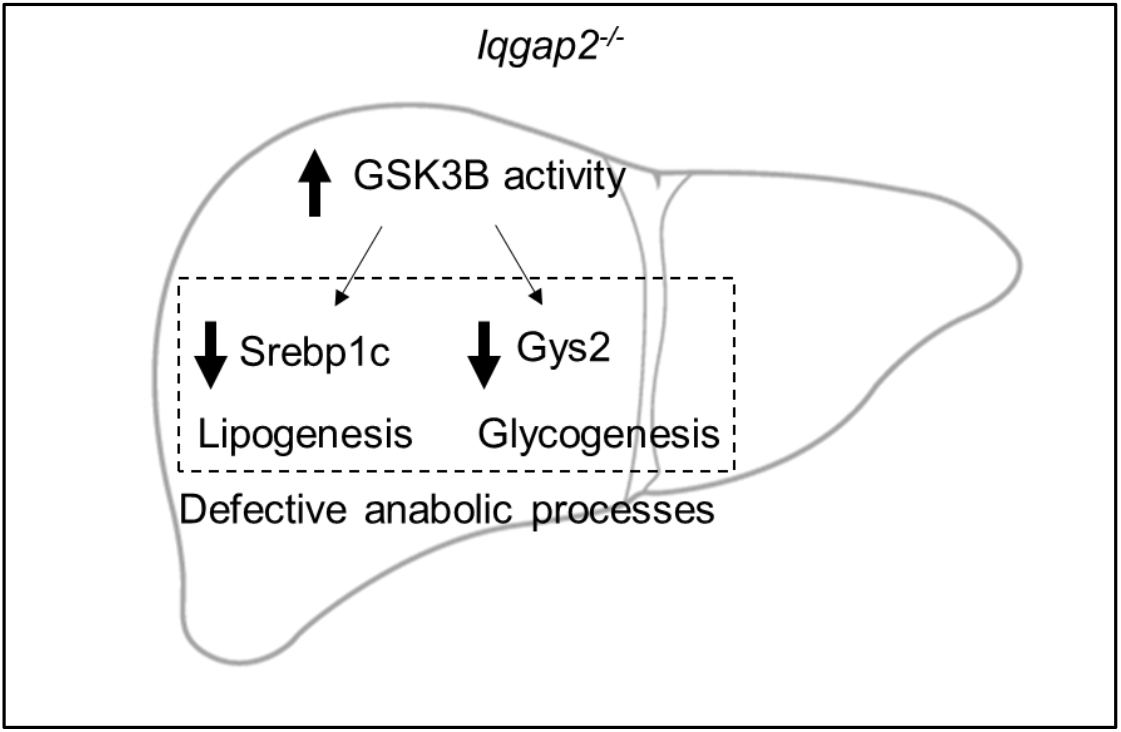
Schematic depicting IQGAP2’s role in regulating GSK activity and liver metabolism.

Since the loss of glycogen in *Iqgap2*^*−/−*^ livers showed zonation pattern, we investigated, protein levels of β-catenin, which not only controls hepatic zonation but is also a downstream target of GSK3 (35, 36, 37, 38). Surprisingly, we found no difference in total β-catenin protein levels in *Iqgap2*^*−/−*^ livers (Fig. 5C, D). It has been shown that β-catenin activity can be altered despite no change in total protein levels (39), we compared its activity between *Iqgap2*^*−/−*^ and WT controls. We examined protein expression of glutamine synthetase (GS), a bonafide β-catenin target to measure β-catenin’s activity (Figure 5E). *Iqgap2*^*−/−*^ livers displayed lower intensity of GS staining compared to WT indicating decreased β-catenin activity. Taken together, our findings suggest that in the absence of IQGAP2, GSK3 is more active resulting in reduced glycogen synthase and β-catenin activity. This in turn results in periportal depletion of hepatic glycogen levels in the fed state.

## Discussion

IQGAP2 has previously been implicated in regulating fatty acid uptake and lipogenesis (23, 24). In this study we demonstrate that IQGAP2 plays a broader role in energy homeostasis, regulating several anabolic processes including those involved in lipid synthesis, glycogen synthesis, and gluconeogenic genes in a sex-specific manner.

We examined if IQGAP2 was involved in mediating fed and fasted state responses in the liver. Although *Iqgap2*^*−/−*^ mice displayed a slightly higher body weight at baseline (23, 40) the male animals lost weight to the same extent as the WT males, but females did not upon fasting (7).

Since we previously showed that IQGAP1 is important for prolonged ketogenesis (20), we examined if IQGAP2 had a similar role. Further, two studies had shown contrasting data with FASN expression being increased and decreased upon fasting in *Iqgap2*^*−/−*^ male livers (23,24). It is possible that Vaitheeswaran et al. used IQGAP2 deletion in the C57/B6 strain under the refed state in their analysis which may explain the discrepancy. Our studies showed the downregulation of *Fasn* and *Srebp1c* after 24 hours of fasting. Although this reduction in *Srebp1c* expression is similar between both sexes of *Iqgap2*^*−/−*^, *Fasn* suppression is severe in female *Iqgap2*^*−/−*^ mice.

We then investigated if the protein expression of IQGAP2 is sexually dimorphic. We observed similar expression of IQGAP2 between males and female livers, which ruled out this possibility. However, metabolic differences in fasting response between male and female mice are well-established, with females having a greater capacity to store and utilize hepatic lipids (41). Hence, this corroborates why we see a severe fasting defect in female *Iqgap2*^*−/−*^ mice.

Fed and fasted responses are orchestrated via hormones and enterokines like insulin, glucagon, and FGF21. It was shown that *Iqgap2*^*−/−*^ mice have normal insulin levels (23) and we demonstrate that starvation hormone *Fgf21*, FGF receptor, *Fgfr1*, and adaptor b-klotho mRNA levels are comparably expressed in these animals. It is possible that despite having an intact expression of hormone and receptors, downstream signaling cascades may be affected as IQGAP2 is a scaffold protein (10). Next we considered the possibility that mitochondrial defects could result in the observed metabolic dysregulation. However, expression of *Pgc1-a*, a key gene in mitochondrial biogenesis, as well as mitochondrial complexes, *CoxII* and *CoxIII* expression were unaltered in *Iqgap2*^*−/−*^ mice. Our data was in line with previous data which showed no difference in mitochondrial function and genomic integrity (42).

To explain the difference in the fed/fasting response in *Iqgap2*^*−/−*^ mice, we extensively examined the anabolic processes. During the fed state, glucose transporters in the liver promote glucose uptake and metabolizes it via glycolysis and store the excess as glycogen. Compared to WT mice, IQGAP2 deficiency did not affect the mRNA expression of *Glut1* and *Glut2*, and phosphofructokinase, a committed step of glycolysis.

However, we found that IQGAP2 deletion in female livers resulted in an overall decreasing trend in genes responsible for gluconeogenesis as well as glycogen synthase expression. *Iqgap2*^*−/−*^ male mice did not display these expression changes in carbohydrate metabolism. Upon further examination, we found that the decrease in glycogen synthase expression correlated well with the periportal depletion of hepatic glycogen levels in *Iqgap2*^*−/−*^ fed female mice. Even though, the gene expression was not seen in *Iqgap2*^*−/−*^ male mice, they also displayed visible periportal depletion of hepatic glycogen. Hepatic glycogen depletion may be due to decreased glycogen synthesis or increased glycogenolysis. Previous findings showed decreased glycogenolysis in *Iqgap2*^*−/−*^ mice (24), therefore we postulated that this glycogen depletion is secondary to changes in glycogen synthesis.

Glycogen synthase kinase (GSK3) is a known regulator of glycogen synthesis. Active GSK3 inactivates glycogen synthase by phosphorylation, which results in lower glycogen synthesis (3). Therefore, during the fed state GSK3 is inactivated by phosphorylation at serine-9 to promote glycogen production. We found reduced phosphorylation of GSK3 in *Iqgap2*^*−/−*^ fed mice suggesting that they have increased active GSK3B, which in turn could inactivate glycogen synthase. This finding is in contrast with previous data, which showed increased pGSK3 expression in *Iqgap2*^*−/−*^ mice (23). This discrepancy is because it was studied immediately after an acute insulin injection not mimicking ad libitum fed state. GSK3B can inhibit β-catenin, which is known to regulate hepatic zonation (35, 36). Therefore, we examined the expression of glutamine synthetase, a well-known downstream target of β-catenin. The reduced expression of glutamine synthetase corroborated lower β-catenin activity in *Iqgap2*^*−/−*^ mice.

GSK is constitutively active and is inhibited either by Ser-9 phosphorylation or protein-binding sequestration (43, 44). It has been shown that GSK3B activity can be regulated by scaffolds like AKAP220 (45). Further studies will be needed to decipher if IQGAP2 can interact directly or via AKAP220 to regulate GSK3 phosphorylation and activity. GSK3 inactivation during the fed state is crucial in coordinating lipogenic and glycogenic metabolism (46, 47). Consistent with this, *Iqgap2*^*−/−*^ mice displayed periportal glycogen depletion and blunted expression of genes involved in anabolic pathways (glycogenesis and lipogenesis). IQGAP2 deficiency disrupted energy homeostasis, more prominently in females. Our data suggest IQGAP2 can modify GSK3 activity and thus subsequent metabolic processes, highlighting a novel role for IQGAP2 in glycogen metabolism.

## Experimental procedures

### Animal Experiments

Generation of *Iqgap2*^*−/−*^ mice has been previously described (42). WT (SVJ/129) and *Iqgap2*^*−/−*^ mice were obtained from Valentina Schmidt, Stony Brook University, Stony Brook, New York, USA. The mice were maintained on a 129/SVJ background and were housed in flow cages at 24°C on a 12/12-hour-light/dark cycle, with lights on starting at 6 AM CST, corresponding to zeitgeber time (ZT) 0. Female and male 12-16-week-old mice were used for all experiments. WT and *Iqgap2*^*−/−*^ mice were randomly divided into ad libitum fed group or a 24-hr fasting group. Fed mice were placed on a normal chow diet (Teklad F6 Rodent Diet, 8664) and allowed ad libitum access to food and water. For the 24-hr fasting experiments, mice were placed in new cages without food, initiating fasting at 9 AM (ZT4) before the day of sacrifice. All mice were sacrificed at ZT4-6. This study was not blinded. Genotype was confirmed by PCR analysis of genomic DNA (n = 4-10 mice per genotype) as previously described (48). A portion of each tissue was fixed in 10% formalin for histological analysis.

### Histology

Formalin-fixed liver sections were stained to assess general histology (Hematoxylin and Eosin), glycogen content (Periodic Acid Schiff Stain (PAS), and PAS/D (PAS with Diastase)), and GS (glutamine synthetase) intensity levels. Methods for each stain are described in detail below.

### Hematoxylin and Eosin Stain

Formalin-fixed liver samples were embedded in paraffin wax, and five-micron sections were cut. The tissue samples were deparaffinized with xylene (3 times for 5 min) and hydrated through a series of decreasing concentrations of ethanol (3 washes of 100% ethanol, followed by 95%, 80%, and 50% ethanol for 3 min each). The samples were then placed in distilled water for 1 min before stained using Hematoxylin 7211 (Richard-Allan Scientific) for 2 min and bluing reagent (0.1% sodium bicarbonate) for 1 min. Slides were placed under running DI water in between both steps until clear. Slides were then placed in 95% ethanol for 2 min before being placed into Eosin-Y (Richard-Allan Scientific) for 25s. The samples were dehydrated in 3 washes of 100% ethanol before being cleared in xylene. Samples were left in xylene overnight. Coverslips were mounted using Permount mounting media (Fisher Chemical). Images were taken using EVOS Microscope (AMG) at 20x magnification, using Bright-Field (BF) microscopy.

### Periodic Acid Schiff (PAS) Stain

Formalin-fixed liver samples were deparaffinized and hydrated to water as listed above. For each PAS-labeled sample slide, there was a corresponding PAS/Diastase (PAS/D)-labeled sample slide. Diastase is an enzyme that breaks down glycogen and PAS/D samples were used as controls. The PAS/D-labeled samples were immersed in distilled water for 2 min before being placed into a 50°C water bath with the diastase (with a-amylase) solution. The diastase solution was made by combining 0.05g diastase (MP Biomedicals) with 50mL of phosphate buffer (0.35g sodium phosphate, dibasic (Na_2_HPO_4_); 3.5g sodium phosphate, monobasic (NaH_2_PO_4_); 200mL water; and pH adjusted to 5.8). PAS/D-labeled slides were left in diastase solution for one hour, while their corresponding PAS-labeled slides were left in distilled water for the same amount of time. The PAS/D-labeled slides were then rinsed in running tap water and immersed in distilled water for 2 min. All slides were oxidized in 0.5% periodic acid solution (0.5g periodic acid (Sigma-Aldrich) in 100mL of distilled water) for 5 min and rinsed in distilled water. Slides were placed in Schiff Reagent (Acros Organics; Catalog: 611175000) for 15 min in a foil-covered container and washed in lukewarm tap water for 5 min. Sections transitioned from a light pink to dark pink color at this stage, an indication of the Schiff Reagent. Samples were counterstained using Hematoxylin 7211 for 20s and washed in tap water for 5min. Samples were dehydrated in 3 washes of 100% ethanol before being cleared in xylene. Samples were left in xylene overnight. Coverslips were mounted using Permount mounting media. Images were taken using EVOS Microscope (AMG) at 20x magnification, using Bright-Field (BF) microscopy.

### Immunohistochemical Stain

Formalin-fixed liver samples were deparaffinized and hydrated to water, as described above. Slides were fully submerged in antigen retrieval sodium citrate buffer (10 mM sodium citrate; 0.05% Tween 20; pH 6.0) and heated in a microwave for 30 min. Quenching using endogenous peroxidase for DAB staining was performed by incubating slides in 3% hydrogen peroxide (Macron Fine Chemicals) with methanol for 15 min in 4°C. Slides were washed in 1X TBST (0.025%) for 5 min before and after the quenching step. After incubating slides with blocking buffer (10% Normal Goat Serum; 1% BSA; 0.1% Tween 20; 0.05% Triton X-100 in 1X TBS) for 1 hour in a humidity chamber at room temperature, primary antibody (1:500 with dilution buffer (1% BSA; 0.05% Triton X-100 in 1X TBS)) was added. Slides were incubated in the humidity chamber in 4°C overnight. Secondary antibody (1:500 with dilution buffer) was added, and slides were incubated in a humidity chamber for one hour at room temperature. Slides were washed with 1X TBST at intermediate steps. DAB reagent from the DAB peroxidase substrate kit (Vector Laboratories) was added per section based on manufacturer’s protocol and the reaction was halted after 4 min by submersion into tap water. After rinsing slides in running tap water for 5 min, slides were counterstained with hematoxylin for 15s. Slides were then submerged in bluing reagent for 30s. Slides were rinsed with tap water in between steps. The samples were dehydrated in one wash of 95% ethanol for 3 min, followed by 3 washes of 100% ethanol for 5 min, before being cleared in xylene. Samples were left in xylene overnight. Coverslips were mounted using Permount mounting media. DAB reagent and TBST were disposed appropriately. Images were taken using EVOS Microscope (AMG) at 20x magnification, using Bright-Field (BF) microscopy.

### Western Blot Analysis

Protein was isolated and homogenized from frozen whole liver tissue through sonification in RIPA buffer (25mm Tris pH 7-8; 150mm NaCl; 0.5% NaDeoxycholate; 1% Triton X-100; 0.1% SDS; 0.1% phosphatase inhibitor; 1 tablet protease inhibitor; in water). Supernatant was collected, and protein concentration was measured using a BCA protein assay kit (Thermo Scientific). For western blots, 30-50μg of sample protein was run through a 5% stacking gel (40% acrylamide; 1.5M Tris pH 6.8; 10% SDS; 10% APS; water; and TEMED) and then 8% resolving gel (40% acrylamide; 1.5M Tris pH 8.8; 10% SDS; 10% APS; water; and TEMED) at 70V for 3 hours. Proteins were transferred from the gel to a PVDF membrane at 35V, overnight at 4°C. Membranes were stained with Ponceau S, rinsed with water, and imaged for the detection of protein bands. Afterward, membranes were washed with 1X TBST and blocked with blocking buffer (5% milk in TBST) for 1 hour on the shaker. Membranes were then incubated in primary antibodies (5% BSA buffer or 5% milk in TBST) overnight at 4°C on the shaker. Secondary antibodies (5% milk in TBST) were added, and membranes were incubated for 1 hour on the shaker at room temperature. Membranes were washed with TBST between steps. Detection of target proteins was performed using SuperSignal West Femto Luminol/Enhancer solution and SuperSignal West Femto Peroxide Buffer (Thermo Scientific). Information about primary and secondary antibodies used are listed in Table S1.

### Quantitative Real time-PCR Analysis

RNA from frozen whole liver tissue was isolated using TRIzol solution (Ambion) based on the manufacturer’s protocol. RNA quality was determined using A260/280 and bleach RNA gel as previously described (49). RNA (3 μg) was treated with DNase (New England Biolabs) and reverse transcribed using random primer mix (New England Biolabs) and the Maxima Reverse Transcriptase kit (Thermo Fisher Scientific). The cDNA was diluted to 12.5 ng/μl with molecular-grade water (Corning) and used for qRT-PCR assays. qRT-PCR was performed on an Eco Real-Time PCR system (Illumina) in triplicates using PerfeCTa SYBR Green FastMix (Quanta). All assays were run with an initial activation step for 10 minutes at 95°C, followed by 40 cycles of 95°C for 15 seconds and 60°C for one minute. *Tbp* was used as the housekeeping gene. Primer sequences of genes used are listed in Table S2. Complete unedited blots are shown in Fig. S6.

### Statistical Analysis

All statistical analyses were performed using GraphPad Prism software. Two-way ANOVA with Bonferroni multiple comparisons test was performed to compare two groups with two treatments. Significance was determined by P < 0.05. Outliers were determined using Grubbs’ test and removed from analysis.

### Study Approval

All animal studies were approved by the University of Illinois at Urbana-Champaign Institutional Animal Care and Use Committee.

## Data availability

All data provided are contained within this manuscript.

## Acknowledgements

We would like to thank Valentina Schmidt (Stony Brook University, Stony Brook, New York, USA) for sharing the *Iqgap2*^*−/−*^ mice. We also thank Hanna Erickson for initial fasting experiments and sharing control samples.

## Supporting information

**Fig. S1.**
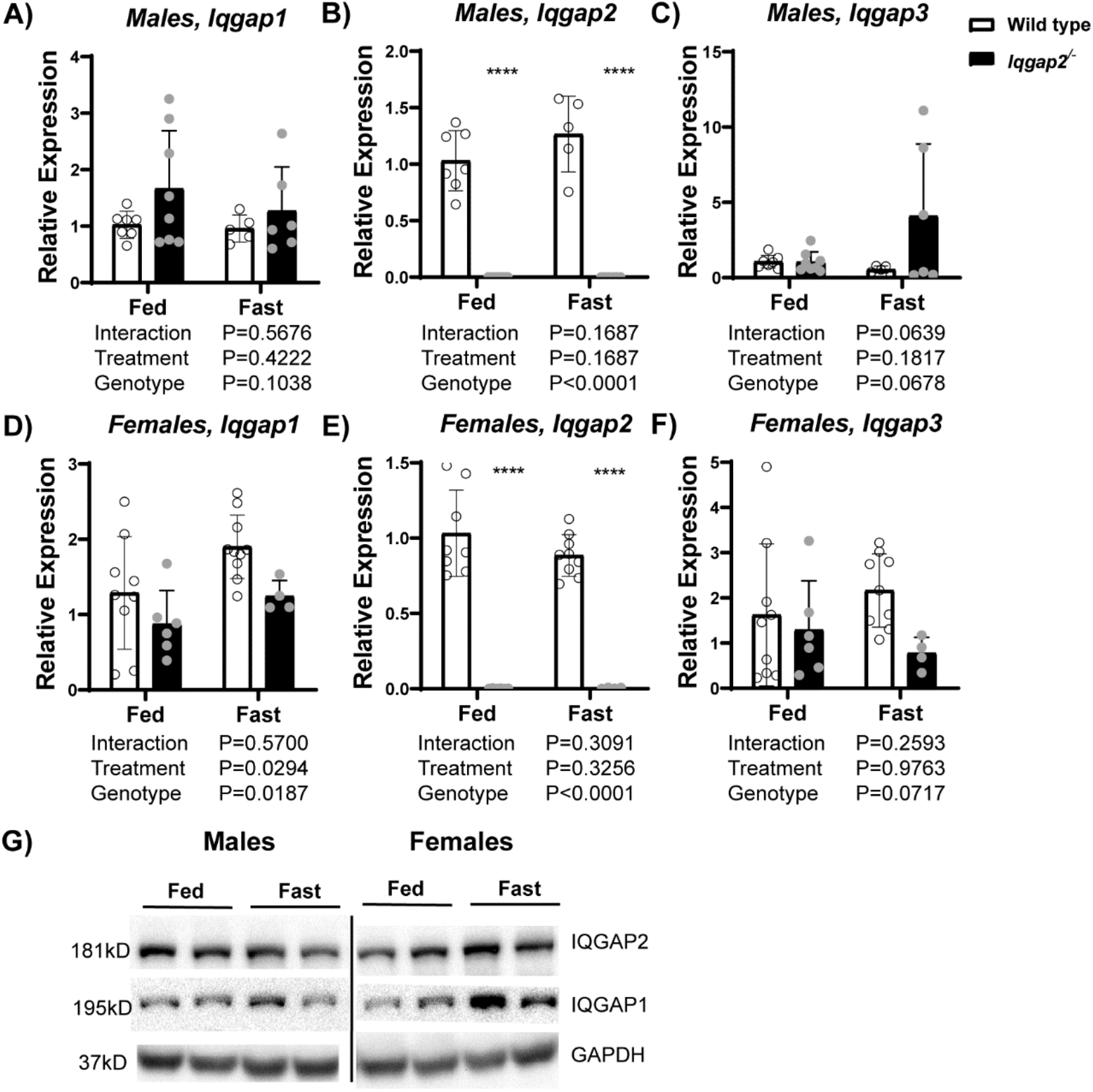
Expression of IQGAPs in Iqgap2^−/−^ mice. We fed regular chow ad libitum or fasted *Iqgap2*^*−/−*^ and WT mice (males and females, 14-16wks old, n=5-15) for 24 hours. Liver tissue was harvested and hepatic expression of IQGAP1 and IQGAP2 at mRNA and protein levels was determined using real-time qRT-PCR and western blots. Absence of IQGAP2 did not alter *Iqgap1*(A,D) and *Iqgap3* transcript expression (C,F) and fasting did not induce *Iqgap2* transcript expression (B, E). Similar expression of IQGAP2 and IQGAP1 protein was noted in both sexes (G), with an induction of IQGAP1 in female fasting state. Statistics were calculated using two-way ANOVA with Bonferroni post hoc analysis. ****P<0.0001 vs WT control.

**Fig. S2.**
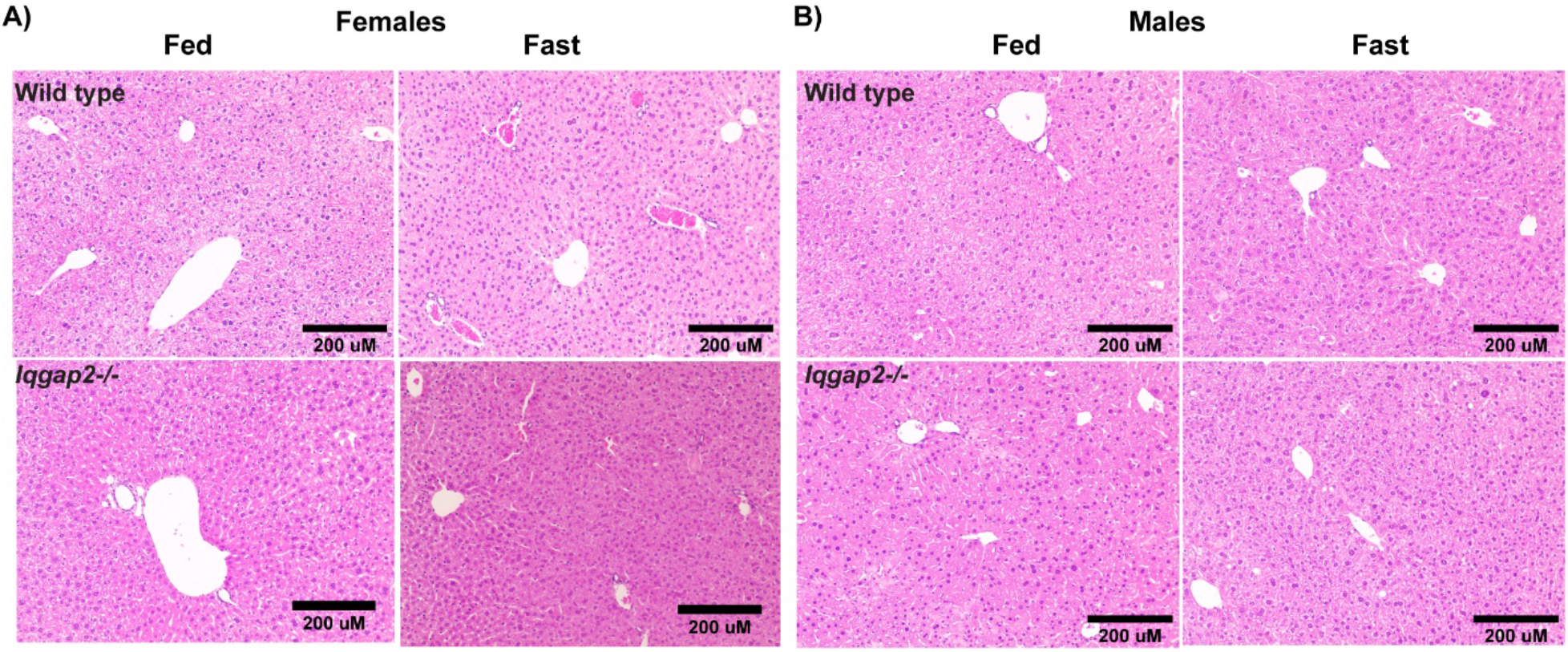
Histological analysis reveals unaltered liver architecture in Iqgap2^−/−^ mice. Liver tissues from *Iqgap2*^*−/−*^ and WT mice were sectioned and stained using hematoxylin and eosin. We do not observe any overt inflammation or changes in hepatocyte size in both (A) female and (B) male livers. N=3-5 per group. Bar, 200μM.

**Fig. S3.**
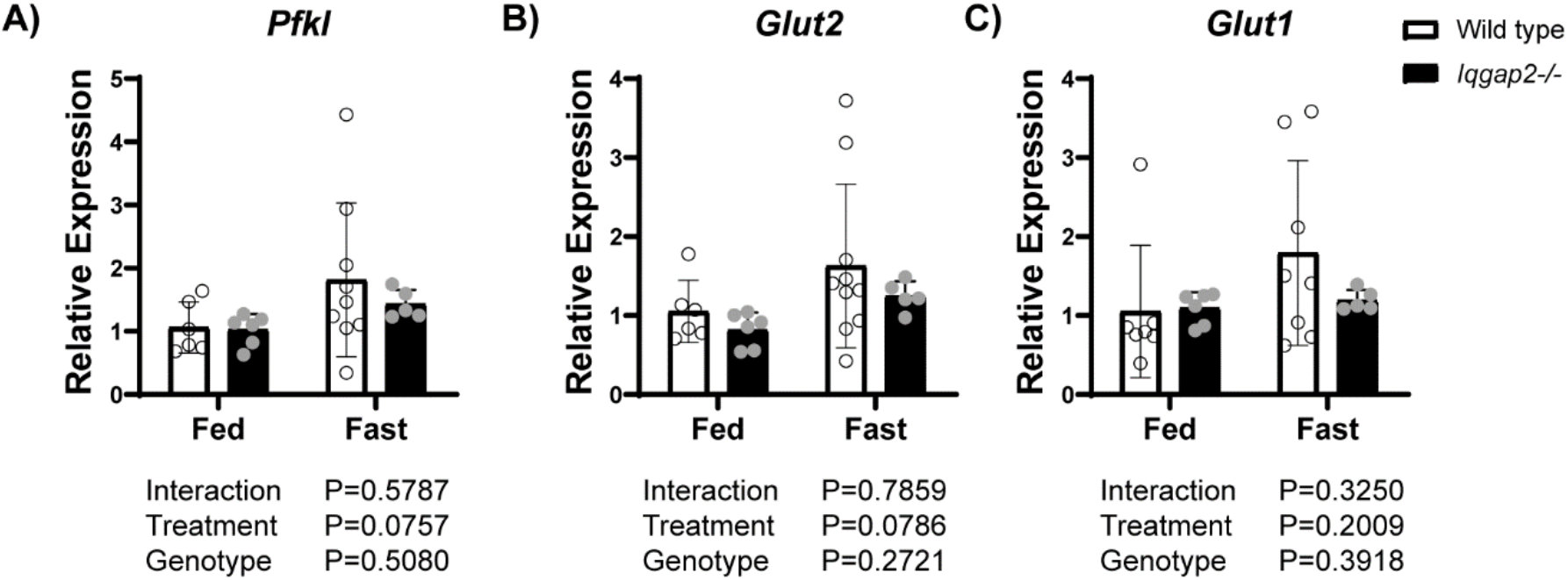
Iqgap2^−/−^ mice do not exhibit any change in glucose uptake and glycolysis. Liver tissues from female *Iqgap2*^*−/−*^ and WT mice were analyzed for the mRNA expression of key genes involved in glucose metabolism. *Iqgap2*^*−/−*^ mice showed no difference in the expression of glycolysis regulator (A) phosphofructokinase and glucose uptake transporters (B, C). Statistics were calculated using two-way ANOVA with Bonferroni post hoc analysis. N= 5-9 mice per group.

**Fig. S4.**
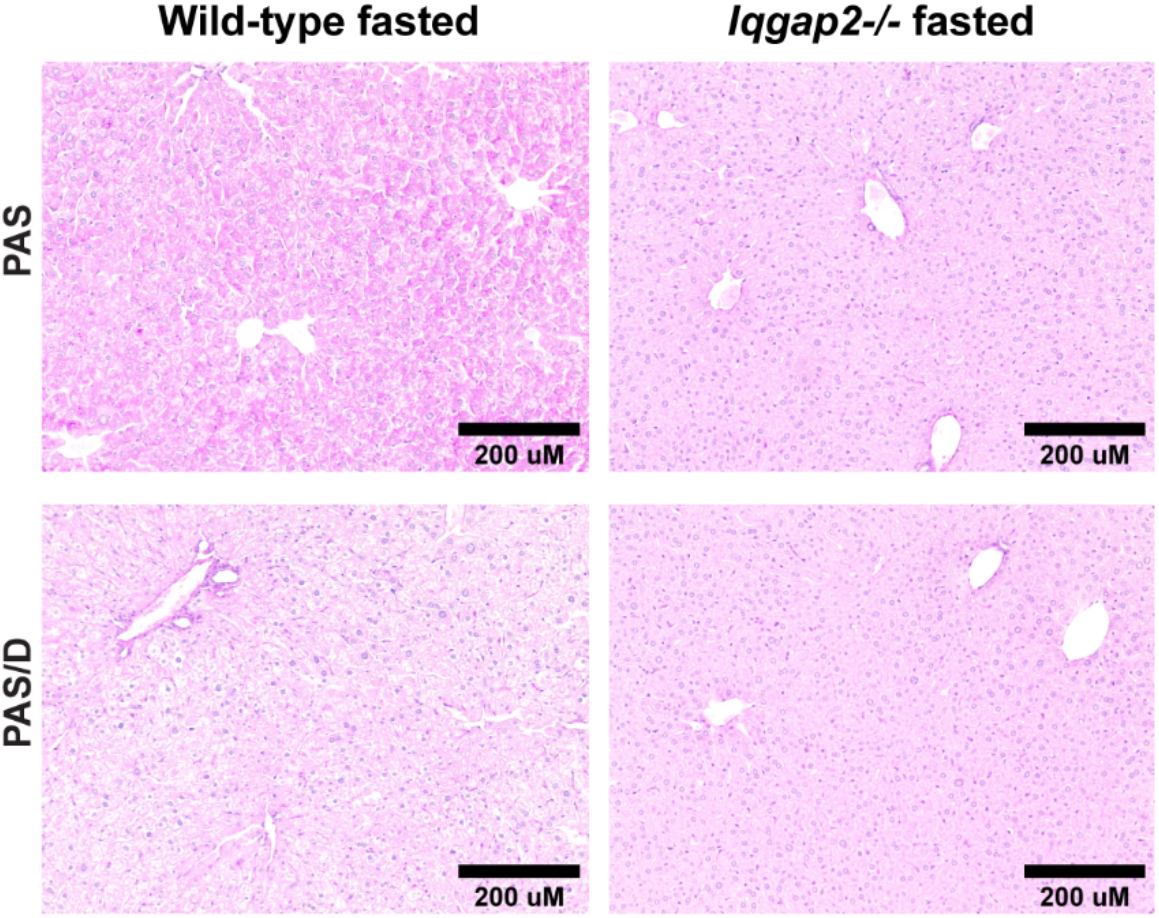
Female Iqgap2^−/−^ mice exhibit intact glycogen mobilization upon fasting. Female *Iqgap2*^*−/−*^ and WT mice were fasted for 24 hours. Liver tissue was histologically stained with Periodic Acid-Schiff’s base (PAS) to examine for glycogen levels. Hepatic glycogen was completely mobilized in both *Iqgap2*^*−/−*^ and WT mice upon 24hours fasting. Bar, 200μM.

**Fig. S5.**
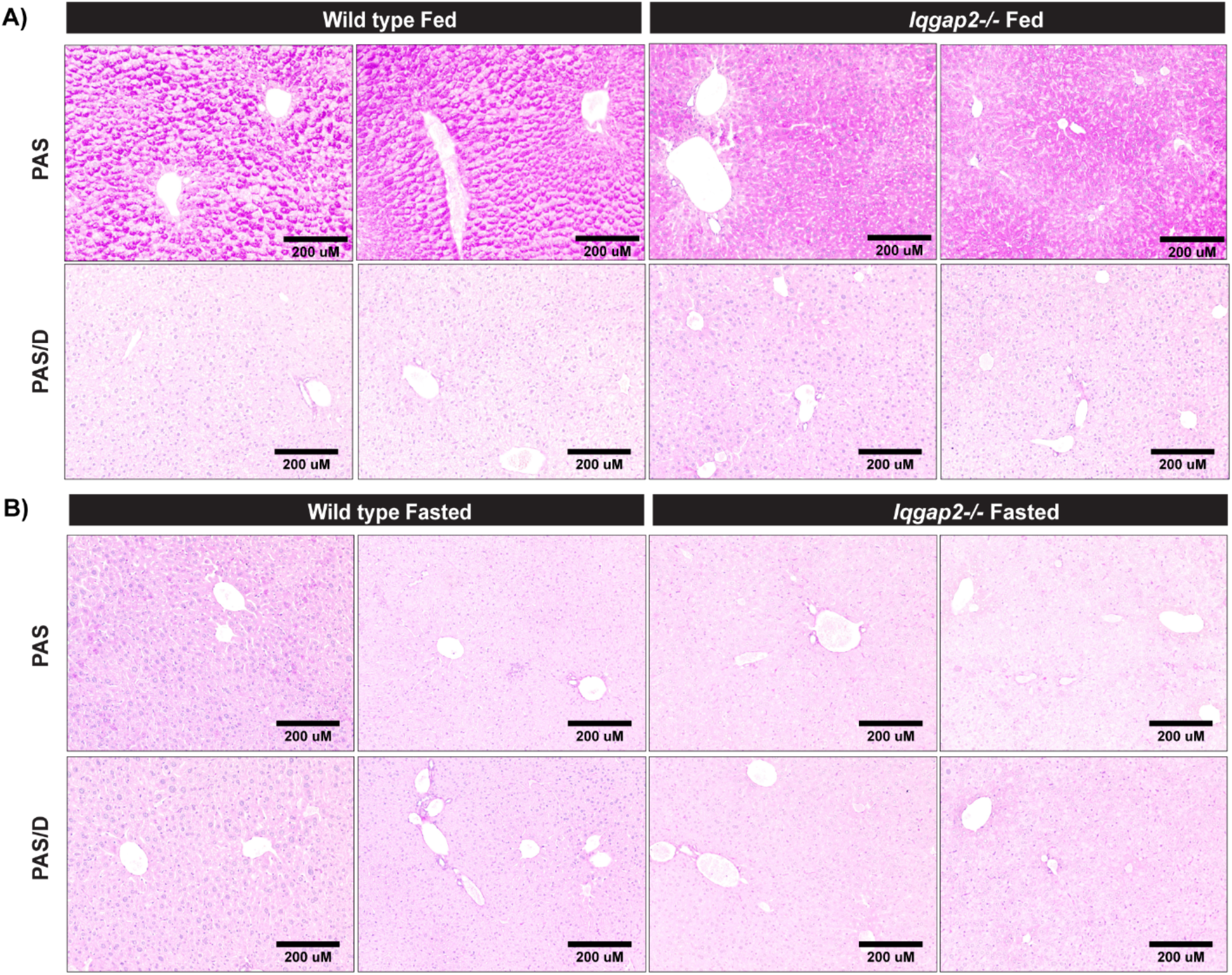
Male Iqgap2^−/−^ mice exhibit lower hepatic glycogen storage. Male *Iqgap2*^*−/−*^ and WT mice were fed regular chow ad libitum. Liver tissue was histologically stained with Periodic Acid-Schiff’s base to (PAS) examine for glycogen levels. Hepatic glycogen in (A) fed and (B) fasted mice was uniformly distributed with a modest decrease of glycogen storage around periportal region in *Iqgap2*^*−/−*^ mice. N=3-4 mice per group. Bar, 200μM.

**Table S1.** Antibodies used for western blots and immunohistochemistry.

| **Antibody** | **Source** | **Catalog number** | **Dilution** | **RRID** |
| --- | --- | --- | --- | --- |
| *Primary antibodies* |  |  |  |  |
| IQGAP2 Rabbit mAb | Abcam | ab181127 | 1:1000 | AB_2833247 |
| IQGAP1 antibody<br>[EPR5220] | Abcam | ab133490 | 1:1000 | AB_11157518 |
| GSK-3B (D5C5Z) XP<br>Rabbit mAb | Cell<br>Signaling<br>Technology | #12456 | 1:1000 | AB_2636978 |
| Phospho-GSK-3a/B<br>(Ser21/9) (D17D2)<br>Rabbit mAb | Cell<br>Signaling<br>Technology | #8566 | 1:1000 | AB_10860069 |
| Glutamine Synthetase<br>Mouse Ab | Fisher<br>Scientific | BDB610518 | 1:500 | AB_397880 |
| Beta-catenin Rabbit Ab | Cell<br>Signaling<br>Technology | #9562S | 1:1000 | AB_331149 |
| GAPDH Rabbit | Sigma | G9545 | 1:5000 | AB_796208 |
| *Secondary antibodies* |  |  |  |  |
| Goat anti-Mouse | ThermoFisher<br>Scientific | #35503 | 1:5000 | 1965946 |
| Goat anti-Rabbit | ThermoFisher<br>Scientific | #31460 | 1:5000 | AB_228341 |

**Table S2.**
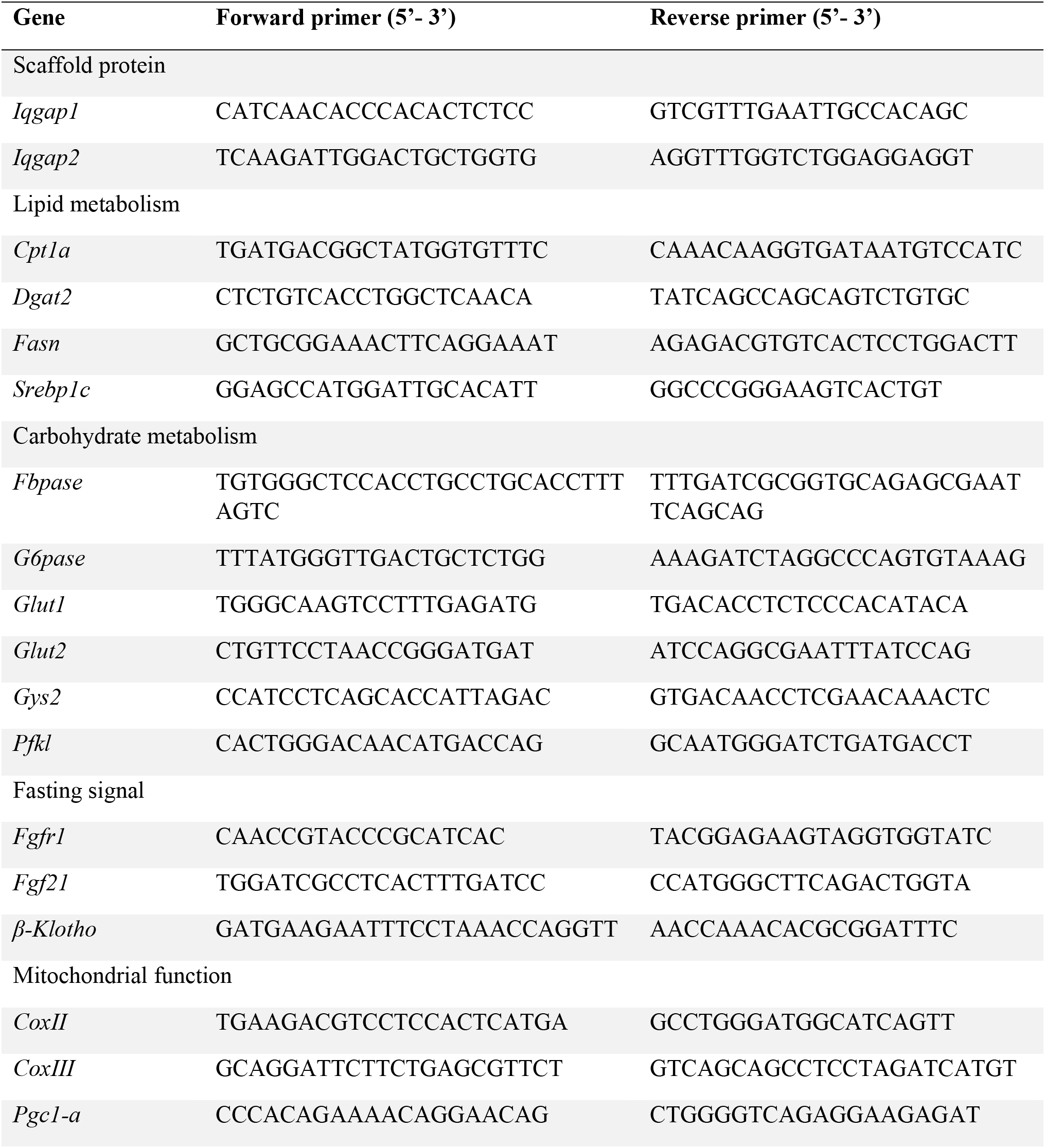
Primer sequences used for real-time qRT-PCR.

**Fig. S6.**
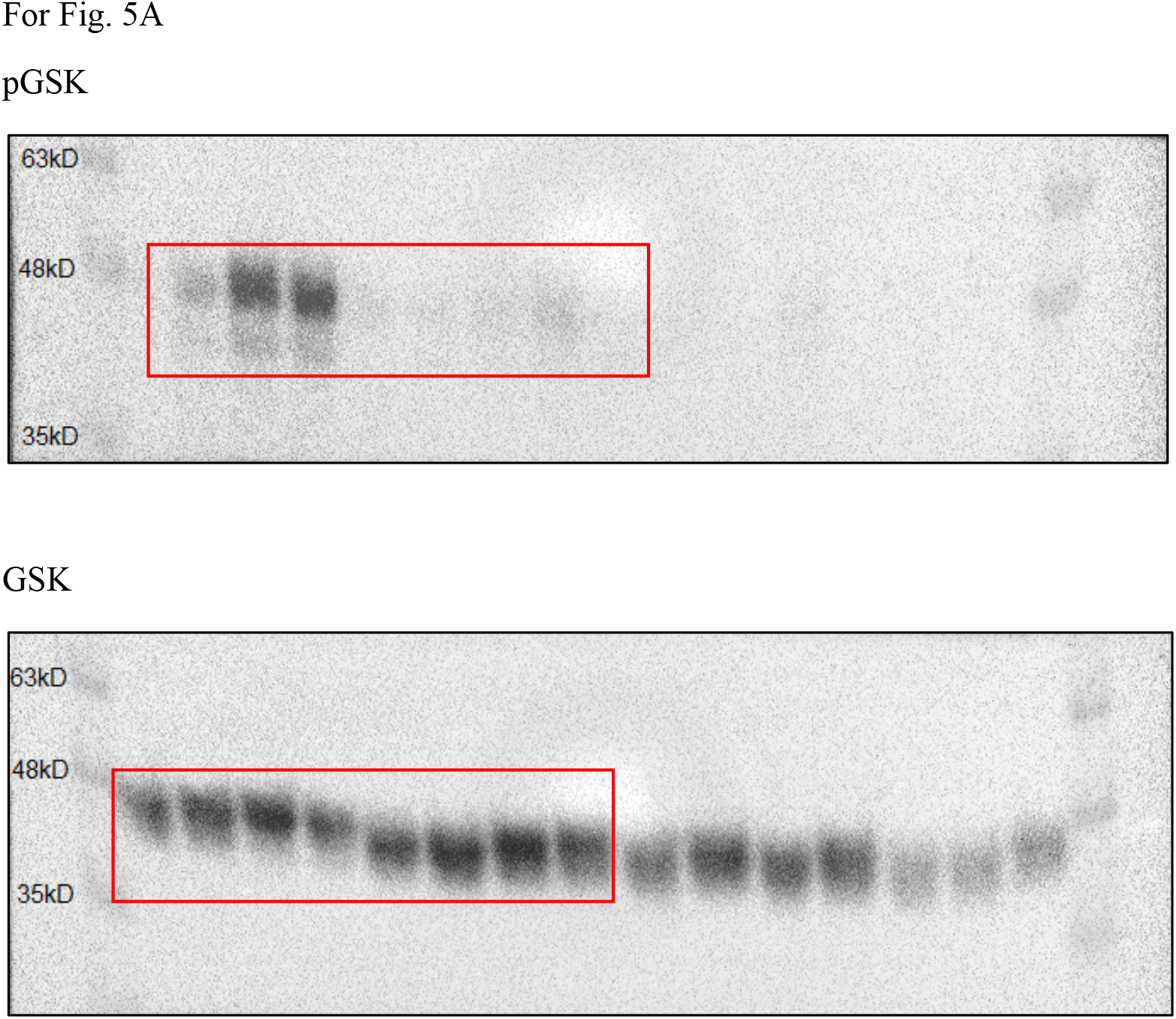
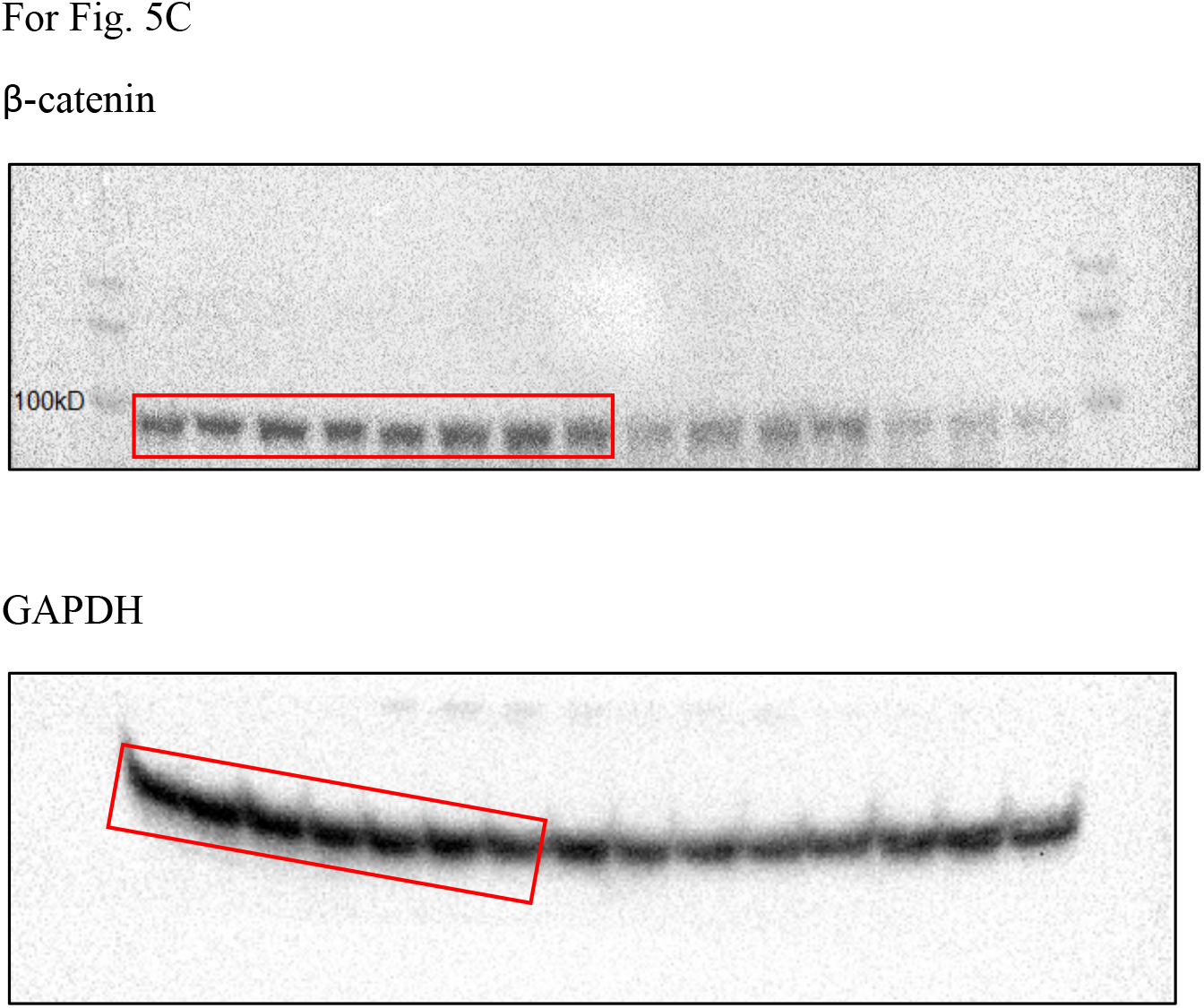
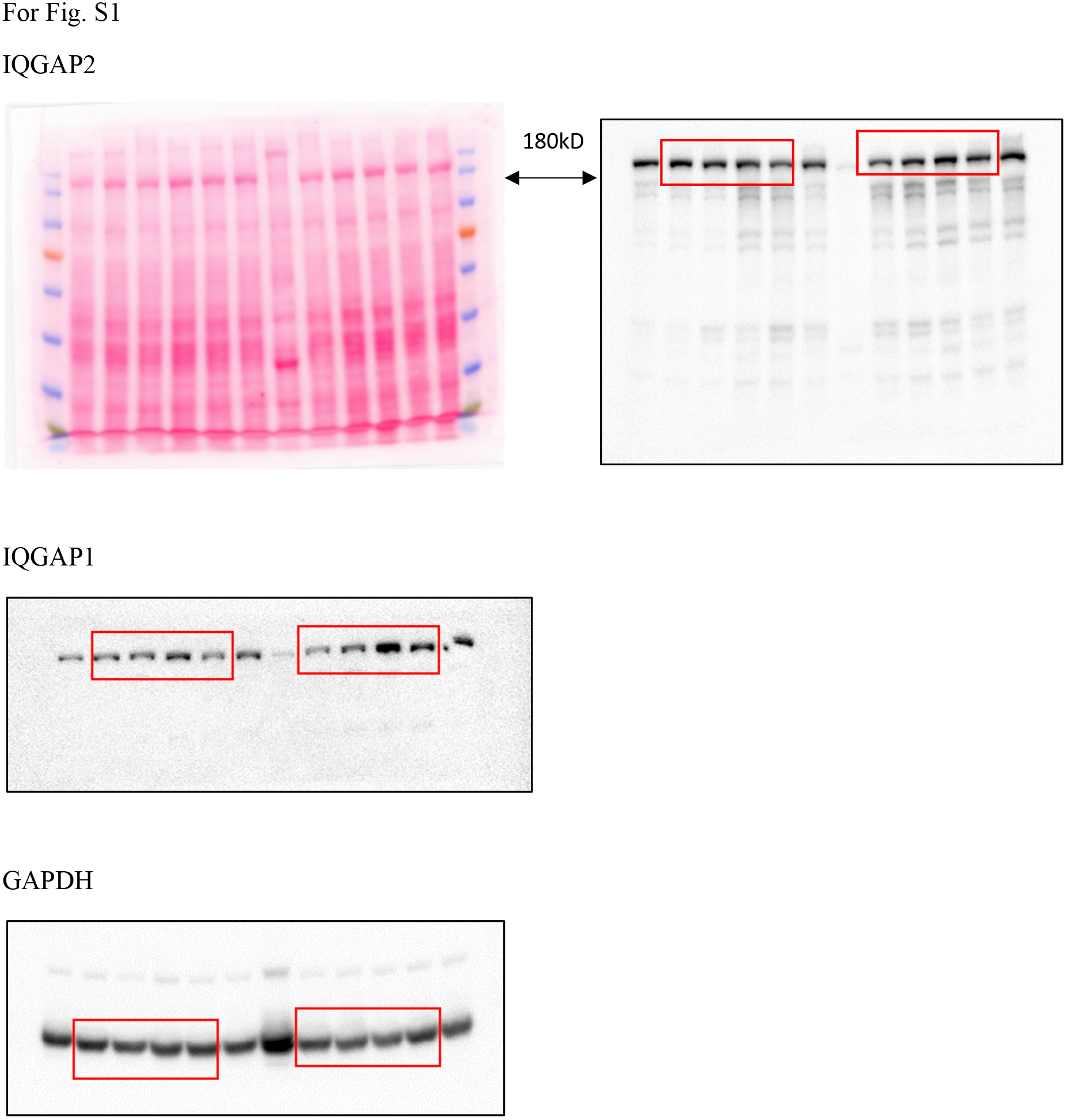
Full unedited gels.

